# Prior unsupervised experience leads to long lasting effects in sensory gating during discrimination learning

**DOI:** 10.1101/2022.09.30.510296

**Authors:** Livia de Hoz, Dana Barniv, Israel Nelken

## Abstract

As the animal moves in its environment, the brain detects and learns the structure of the surrounding stimuli, independently of the immediate relevance this has for the animal. This experience influences subsequent learning in a manner quantified using paradigms such as latent inhibition or stimulus preconditioning, which measure the effect that unsupervised (not-reinforced) learning has on subsequent reinforced learning. Despite our understanding of the behavioural consequences of prior neutral experience, there is little understanding about the influence of this previous experience on neuronal plasticity. Using latent inhibition, we have shown in mice that learning a two tone discrimination is slower in mice that have had previous neutral exposure to the same or similar tones (<2/3 octave away). Neutral exposure thus elicits profound changes in the brain that influence subsequent learning. To study how previous experience influences experience-dependent plasticity, and better understand the interactions between experience, learning, and plasticity, we recorded sound evoked responses in the auditory cortex of exposed and trained mice. We studied both changes in response magnitude and changes in sensory dynamics, and related both to the differential behavioral effects of different pre-exposure conditions. Here we describe the neuronal changes that paralleled the behavioral findings. We found that discrimination learning led to stronger initial sound-evoked responses and a long-lasting increase in response adaptation and an increase. The first effect was delayed in animals that showed latent inhibition, paralleling behavioural learning. Overall our data reveal that slow changes in behaviour that accompanied learning, paralleled the slow dynamics of experience-dependent plasticity in auditory cortex.

## Introduction

Learning in natural environments is a cumulative process that builds upon previous experiences and continuously adapts to new stimulus contingencies. There is considerable behavioral evidence for the influence of previous experience on an animal’s subsequent interpretation of the world, using paradigms such as generalization (Jaramillo, Borges, and Zador 2014; Chen, Krueger-Burg, and de Hoz 2019; Chen et al. 2021), latent inhibition (Lubow 1973; de Hoz and Nelken 2014; Holland 2018) or stimulus preconditioning (Brogden 1939; Rizley and Rescorla 1972; Escobar, Arcediano, and Miller 2002; Headley and Weinberger 2015; Robinson et al. 2014). And yet, there is little understanding about the influence of this previous experience on neuronal plasticity. The aim of this study is to better understand the interactions between experience, learning, and plasticity.

That learning is associated with neuronal plasticity has been shown across modalities and paradigms. In the auditory system, for example, changes in neuronal receptive fields develop in a stimulus-specific manner after associative learning (Bakin and Weinberger 1990; Blake et al. 2006; Polley, Steinberg, and Merzenich 2006; Jaramillo, Borges, and Zador 2014; de Hoz et al. 2018; Dalmay et al. 2019; Ceballo et al. 2019). In addition to changes in sensory coding, learning may modify more abstract properties of cortical responses to sounds. For example, we have shown that fear conditioning modifies stimulus-specific adaptation (SSA; (Yaron et al. 2020)). SSA is the decrease in neuronal responses to a repeated stimulus (adaptation) that does not, or only partially, generalize to other stimuli (Ulanovsky, Las, and Nelken 2003; Taaseh, Yaron, and Nelken 2011; Hershenhoren et al. 2014; Carbajal and Malmierca 2018). SSA may be a mechanism underlying some forms of sensory gating, leading to suppression of information about repeated sounds while keeping sensitivity to changes in sound identity. In this study, we explore how prior experience with sound influences learning-induced changes in sensory dynamics responsible for SSA and therefore for sensory gating.

Using a latent inhibition paradigm, we observed that previous experience influences subsequent behavioral discrimination learning (de Hoz and Nelken 2014). We exposed mice to a sound for a few days in neutral settings, then made it predictive of a negative outcome in an operant learning paradigm. Animals exposed in this manner took longer to learn the new association of the sound with a negative outcome than unexposed animals (latent inhibition). Interestingly, in these paradigms, a change in behavior was found not only with respect to the conditioned tone, but also with respect to other stimuli whose meaning did not change, notably a sound that indicated safe trials. Once animals began to avoid the conditioned tone, they also avoided (at least for a while) reward when safe sounds were played. The generalization to the safe sound took a few days to decline and the decline was slower in animals that showed latent inhibition (de Hoz and Nelken 2014).

Here we describe neuronal changes that paralleled these behavioral findings. To study how previous experience influences experience-dependent plasticity, we recorded sound evoked responses in the auditory cortex (AC) of a subset of the mice trained in de Hoz and Nelken (2014) study. We studied both changes in response magnitude and changes in sensory dynamics, and related both to the differential behavioral effects of different pre-exposure conditions.

## Methods

### Ethics statement and animals

Experiments were performed in The Hebrew University of Jerusalem. The joint ethics committee (IACUC) of the Hebrew University and Hadassah Medical Center approved the study protocol for animal welfare. The Hebrew University is an AAALAC International accredited institute.

We used 42 C57BL/6JOlaHsd female mice obtained from a commercial supplier (Harlan, Israel). The mice were 7–8 weeks old at the time of arrival and were always kept in a light/dark 7am/7pm cycle. A few days after arrival, each mouse was lightly anaesthetized with isoflurane vapour and a sterile transponder (DATAMARS T-IS 8010 FDX-B, 13 mm long, 2 mm in diameter, weight 0.1 gr; or IS0 compliant 11784 transponder, 12 mm long, from TSE) was implanted subcutaneously in the upper back. In later replications, a stitch or tissue adhesive (histoacryl; Braun) was used to close the small hole left on the skin by the transponder injection. Once recovered from anesthesia, the mice were placed in the Audiobox (see below). Seven mice underwent electrophysiological recordings before conditioning (pre-exposure group, see below); the other 35 mice underwent conditioning, and are a subset of the animals used in a previous study that studied latent inhibition and its frequency-dependence (de Hoz and Nelken 2014).

### Apparatus: Audiobox

All behavioral tests were run in an Audiobox (TSE Systems, Germany), a device developed for auditory research and based on the Intellicage (TSE Systems, Germany). The Audiobox serves both as living quarters for the mice and as their testing arena. The mice were kept in groups of 8 to 10 animals. Each animal was individually identifiable through the implanted transponder, and the behavior of each mouse was automatically monitored by two means: reading of the unique transponder by an antenna at the entrance of the drinking corner (see below), and detection of specific behaviors (nose-poking and licking) through other sensors.

The Audiobox was kept in a dedicated temperature-regulated room, used only for these experiments. The Audiobox consists of two compartments connected by a long corridor (Figure 1A). One compartment, a normal mouse cage, served as the home cage, where the animals had access to food ad libitum. Water was delivered in the second compartment of the Audiobox, the ‘corner’ (Figure 1A), positioned inside a sound-attenuated box. Entrance into the corner is termed here a ‘visit’. The beginning of the visit was identified by the detection of the mouse’s transponder by the antenna together with the activation of a heat sensor inside the corner. The stimulus to be presented was selected according to the identity of the mouse as identified by the transponder. Behavioral data were logged for each mouse individually. Once in the corner, the mouse could access water by nose-poking into either of two ports, one at each side of the corner. The doors to the ports were opened or closed depending on the demands of the experiment. A loudspeaker (DSM 25 FFL 8 Ohm from Visaton, or a 22TAF/G from Seas Prestige) was positioned directly behind the corner, slightly above it, for the presentation of the stimuli.

**Figure 1.**
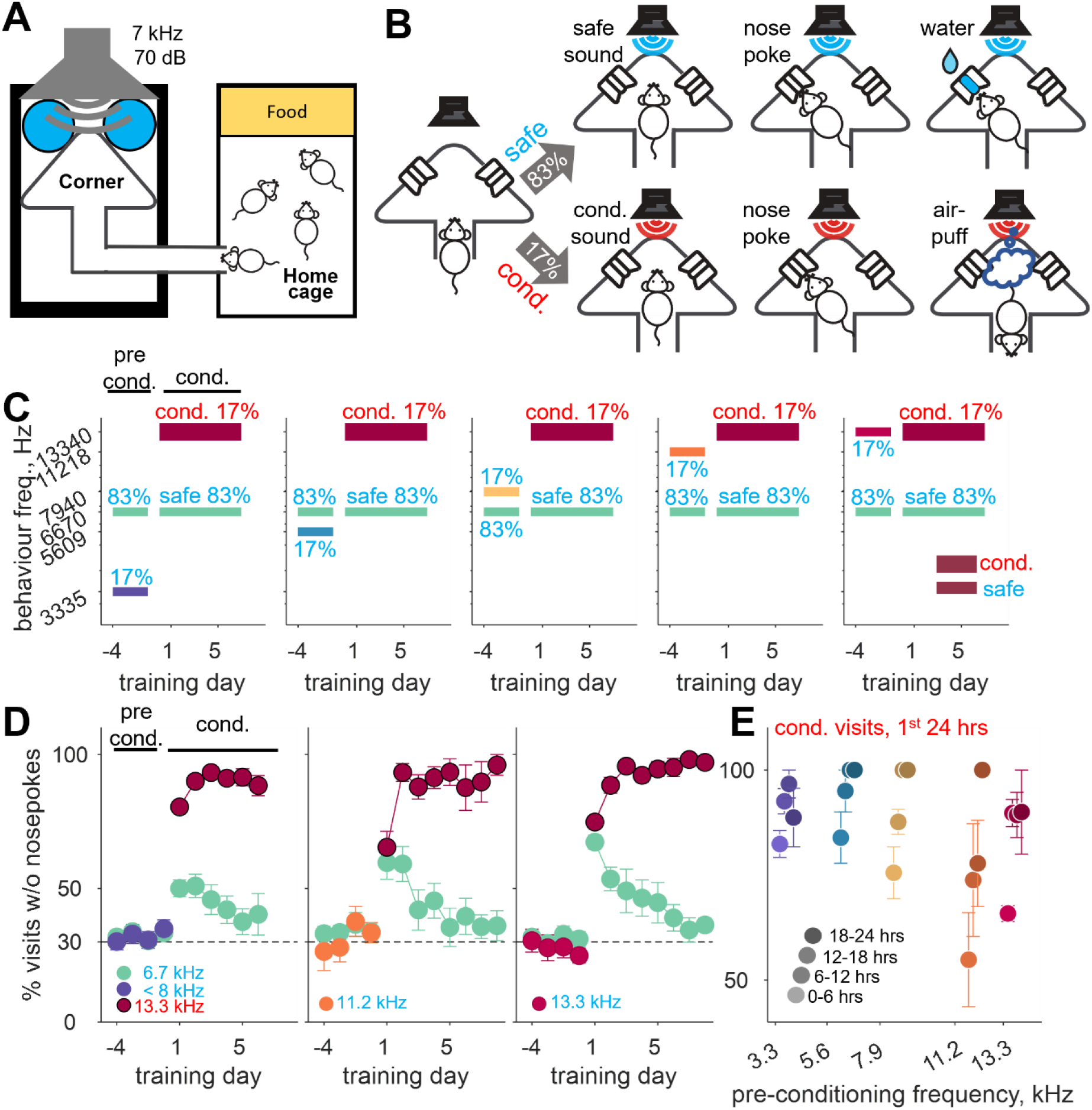
behaviour: latent inhibition in groups exposed to frequencies at or near the conditioned frequency. **A.** Schematic of the Audiobox comprising of a home cage with food ad libitum and a sound attenuated cage containing the water corner. Tones were played for the duration of the visit to the corner. **B.** Diagram explaining ‘safe’ (top) vs ‘conditioned’ (bottom) visits with nose-poke being followed by access to water or air-puff, respectively. **C.** Diagram explaining time course of sound exposure according to groups. All mice were exposed to 2 safe sounds during a pre-conditioning phase (83% visits with 6.7 kHz, 17% of visits with another group-specific frequency). During conditioning all mice had 83% ‘safe’ visits (6.7 kHz) and 17% ‘conditioned’ visits (13.3 kHz). **D.** Behavioral performance by day and pre-exposure frequency (left: < 8 Khz, 18 mice began conditioning; middle: 11.2 kHz, 6 mice began conditioning; right: 13.3 kHz, 11 mice began conditioning). Seven mice were removed from the Audiobox for electrophysiology during preconditioning. Mice nose-poked in about 70% of both types of safe visits during the pre-conditioning exposure phase. When conditioning began, mice learned to avoid nose-poking in conditioned visits (circled red dots) and continued nose-poking in safe visits (6.7 kHz, green). Nose-poking in safe visits was somewhat reduced compared to pre-conditioning, particularly in groups exposed to 11.2 and 13.3 kHz. Dash line: 30% visits without nose-pokes level. **E.** Performance in blocks of 6 hours during the first 24 hours of conditioning colour-coded according to pre-exposure frequency. Mice learned to avoid nose-poking in conditioned visits (13.3 kHz for all mice) but learning was faster in mice exposed to frequencies away from the conditioned sound, below 8 kHz (left, purple, blue and gold). A and B courtesy of Chi Chen.

### Sounds

We used tones with frequencies ranging between 3 and 19 kHz. This frequency range contains the area of highest sensitivity in a mouse audiogram, and include species-specific vocalizations such as pups wriggling calls. Stimuli consisted of 30 ms pure tone pips, with 5 ms rise/fall linear slopes, repeated at a rate of 3 Hz throughout a visit. All tone pips presented in the corner throughout a visit had the same frequency. Sounds were played at a fixed intensity of 70 dB SPL. Sounds were generated using Matlab (Mathworks) at a sampling rate of 96 kHz and written into computer files. The sound level was calibrated using either a Brüel and Kjaer (4939 J0 free field) or a GRAS (1/40 40BE) microphone. The microphone was placed at different positions within the corner, as well as outside the corner. Relevant sounds were played at the nominal intensities used in the study. Microphone signals were sampled at 96 kHz and analyzed in Matlab. Tones between 3 kHz and 19 kHz did not show any significant harmonic distortion. In the rare occasions when harmonics were present, they were at least 40 dB below the main signal. There was a linear correspondence between the nominal sound level and the sound level measured by the microphone. While sounds played inside the corner were significantly attenuated outside of the attenuated box (20 dB), there was little attenuation between the corner and the corridor directly leading to it (about 10 dB). In consequence, mice in the corridor could hear the sound presented to the mouse inside the corner. This, however, did not seem to affect their behavior.

### Behavioral training

The aim of the experiment was to study the effect of prior experience on conditioning. For this purpose, different mice were pre-exposed to different frequencies, at, near, or far from the subsequent conditioned frequency. After the pre-exposure ended, all mice were conditioned to the same conditioned frequency. Latent-inhibition was quantified by the reduction of the efficacy of conditioning during the first few conditioned tone presentations.

Training consisted of the following stages (see Figure 1B):

Habituation phase (7 days): Immediately after the transponder implantation, mice were placed into the Audiobox. During this phase the doors giving access to the water within the corner remained constantly open and no sound was presented during the visits.

Safe-only phase (4 days): Once the mice had learned to access the water and were drinking freely, the doors were closed and only opened when the mouse nose-poked into the port. At the same time, every visit to the corner was coupled, for the duration of the visit, with the presentation of the 6670 Hz safe sound (tone pips).

Pre-conditioning phase (5 days): The pre-exposed tone (whose frequency varied according to the experimental group, see below) was now played in 17% of the visits (randomly once every ~6 visits). A nose-poke in these visits resulted, like in all safe visits, in the opening of the doors and access to water. The remaining 83% visits were associated with the 6670 Hz safe tone as before. Therefore, during this phase, the pre-exposure frequency had the same behavioral significance as the safe frequency. The last 4 days of pre-conditionng were used for the behavioural and electrophysiological analysis presented here.

The pre-exposure tone had one of the following frequencies: 3335 Hz (2 octaves below 13340 Hz, 9 mice), 5609 (1.25 octaves below, 4 mice), 7932 Hz (0.75 octave below, 4 mice), 11218 Hz (0.25 octave below, 6 mice), or 13340 Hz (equal to the conditioned tone, 11 mice). The choice of frequencies was based on ABR threshold measurements (Jackson Laboratories phenotype database) which showed large increase in thresholds between 16 and 32 kHz in this mouse strain. Therefore, we avoided using frequencies above 16 kHz. Generally, the mice in any one replication were divided into 2 or 3 groups that each had a different pre-exposure frequency, and in most replications one of these groups was pre-exposed to the frequency that was conditioned later, 13340 Hz.

Conditioned phase (1-10 days depending on the mice): This phase was identical for all mice, irrespective of their pre-exposure frequency. The preexposure visits were replaced by conditioned visits, also with a probability of 17%. For all mice, the conditioned tone had a frequency of 13340 Hz. During this phase, a nose-poke during conditioned visits resulted in the delivery of an air-puff and the doors were not opened. The remaining 83% of the visits were associated with the 6670 Hz safe tone as before. Therefore, during this phase, the two frequencies that the mouse could hear in the corner had very different behavioral significance.

### Electrophysiology

Mice were removed from the Audiobox one at a time for electrophysiology. The selected mouse was anaesthetized with Avertin (tribromoethanol, 250 mg/Kg body mass, i.p.), and attached to a stereotactic frame using a metal head-bar glued with hystoacryl (Braun) and super glue to the top of the skull, in order to leave the ears free. A metal screw was fixed to the skull and used as ground. A craniotomy over the left auditory cortex was performed after detachment of the temporal muscle. The craniotomy was made following the contour delimited rostrally and ventrally by the temporal fissures, dorsally by the temporal ridge, and caudally by the caudal ridge. It was about 4mm long (anteriorposterior axis) and 2mm wide (dorsoventral axis). A glass coated tungsten electrode (Alphaomega, 900 kOhm impedance) were inserted 350-450 microns deep perpendicular to the cortical surface and both spikes and local field potentials were recorded during sound presentation. Voltage signals were sampled at 25 kHz, pre-amplified (x10), filtered between 3 and 8000 Hz, and amplified again (x5000; MCP, Alphaomega). For offline multiunit spike detection, the signal was band-passed (500–6000 Hz) and negative peaks that exceeded a threshold of 7 times the median of the absolute deviation from the median (MAD) were considered as spikes. Spike time was defined as the time of the peak of the spike.

### Sound stimulation during electrophysiology

All sound generation was performed using Matlab (The Mathworks, Inc.). The digital signals were transduced to voltage signals by a sound card (HDSP 9632, RME), attenuated (PA5, TDT), power-amplified (ED1, TDT) and played through a sealed speaker (EC1, TDT) into the right ear canal of the mouse. For pure tones, 0 dB attenuation corresponded to a sound level of about 100 dB SPL throughout the 3.5-octave range used here (2-24 kHz).

Pure tones of different frequencies (2–24 kHz), 30ms in length and with 5ms rise/fall linear slopes were used to ensure that recordings were in regions with significant responses at 6670 (safe tone) and 13340 Hz (conditioned tone). Oddball paradigms comprising 2 pure tones (ΔF = 1.2 or 1.44%, often 6670 and 13340 Hz) were used to determine SSA. Tones were 30 ms in duration with 5 ms rise/fall linear slopes and were presented at a rate of 3 Hz. Each pair of tones was used in 2 oddball protocols. In one, the lower frequency was presented in 95% of the trials (standard) and the higher tone in 5% (deviant or rare). In the other protocol the lower tone became the rare tone and the higher tone, the standard (Figure 4A). A total of 500 trials was used per oddball paradigm such that the rare and standard tones were presented in 25 and 475 trials, respectively. Each frequency was also presented in a ‘deviant-alone’ block in 25 trials where the remaining 475 were silent.

**Figure 2.**
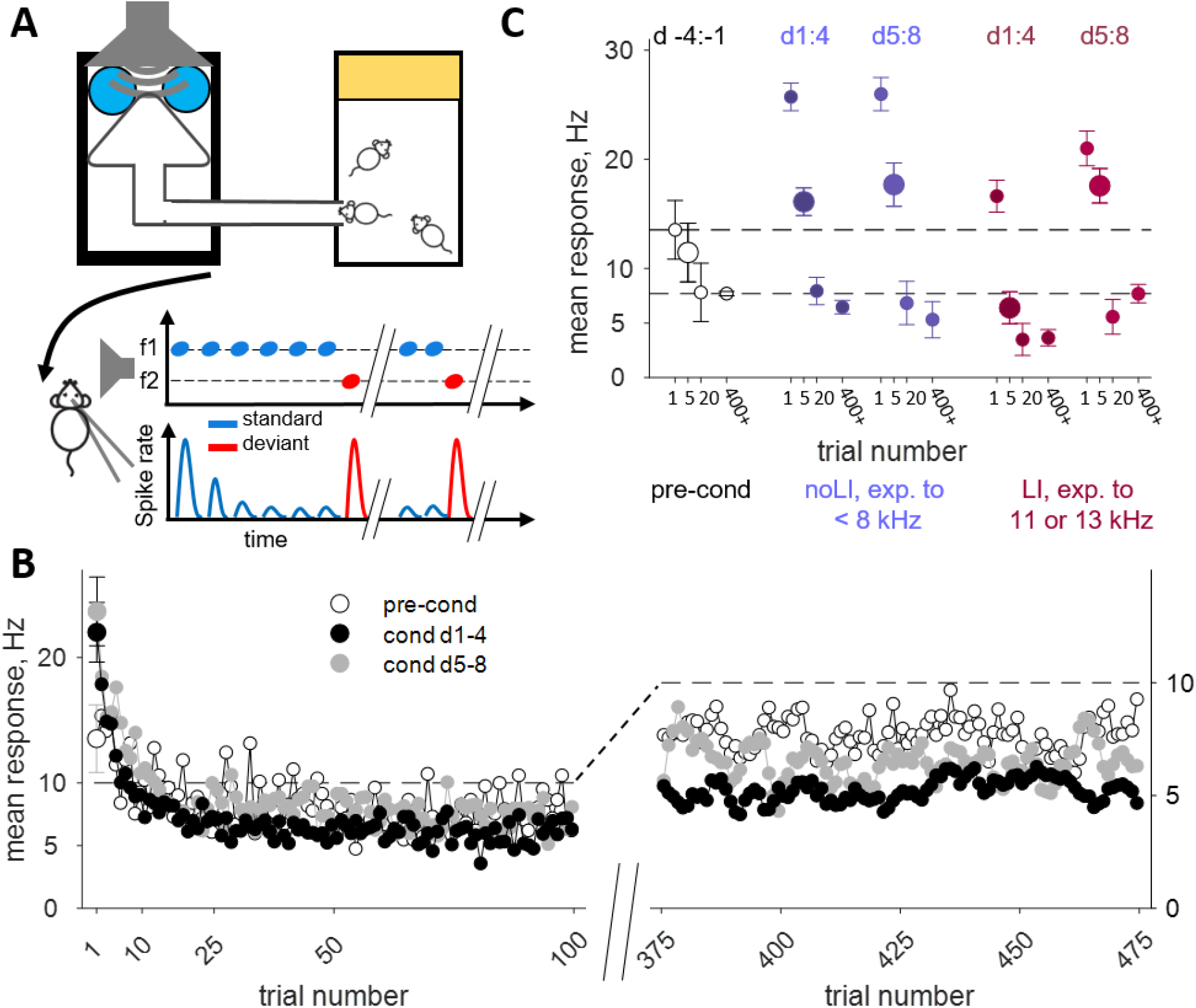
response adaptation was stronger in conditioned mice. **A.** scheme of electrophysiology: one mouse is removed from Audiobox for recordings in A1 during acoustic stimulation in the form of oddball paradigm. One sound (standard, blue) is repeated leading to response adaptation, and one rare sound (deviant, red) evoked non-adapted responses. **B.** Mean response (sum of spikes over 100 ms from stimulus onset) to standard tone across first (left) and last (right) hundred trials across recording sites during preconditioning (white; n=7 mice and 82 recordings), first 4 conditioning days (black; n=18 mice and 212 recordings), and days 5 to 8 of conditioning (gray; n=17 mice and 152 recordings). Error-bars only on trial 1 for clarity. **C.** Mean response to trials 1, 5 (bigger dot for visualization), and 20 as well as mean of last 75 trials according to pre-exposure frequency (color coded) and time-block of conditioning (left: days 1 to 4, or right: days 5 to 8). Preconditioning (white; n=7 mice and 82 recordings), exposed to tones < 8 kHz (purple; conditioning d1-4 n=9 mice and 126 recordings; d5-8 n=9 mice and 82 recordings), exposed to 11.2 or 13.3 kHz (red; conditioning d1-4 n=9 mice and 86 recordings; d5-8 n=8 mice and 70 recordings).

**Figure 3.**
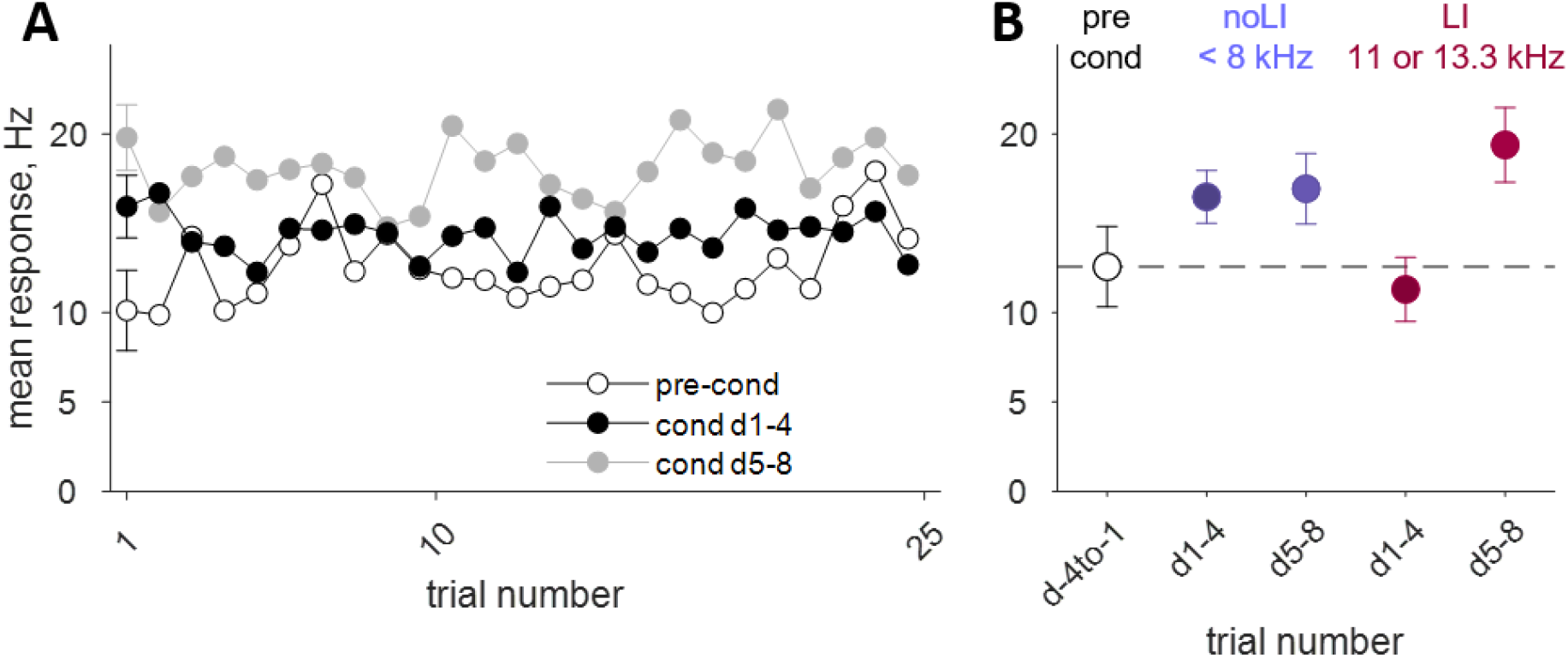
deviant response increases in conditioned mice. **A.** Mean response (sum of spikes over 100 ms from stimulus onset) to deviant tones across all 25 deviant trials across recording sites during preconditioning (white), first 4 conditioning days (black), and days 5 to 8 of conditioning (gray). N values as Figure 2B. **B.** Mean response to all trials according to exposure frequency (color coded) and time-block of conditioning (left dark: days 1 to 4, or right light: days 5 to 8). Preconditioning (white), exposed to tones < 8 kHz (purple), exposed to 11.2 or 13.3 kHz (red).

**Figure 4.**
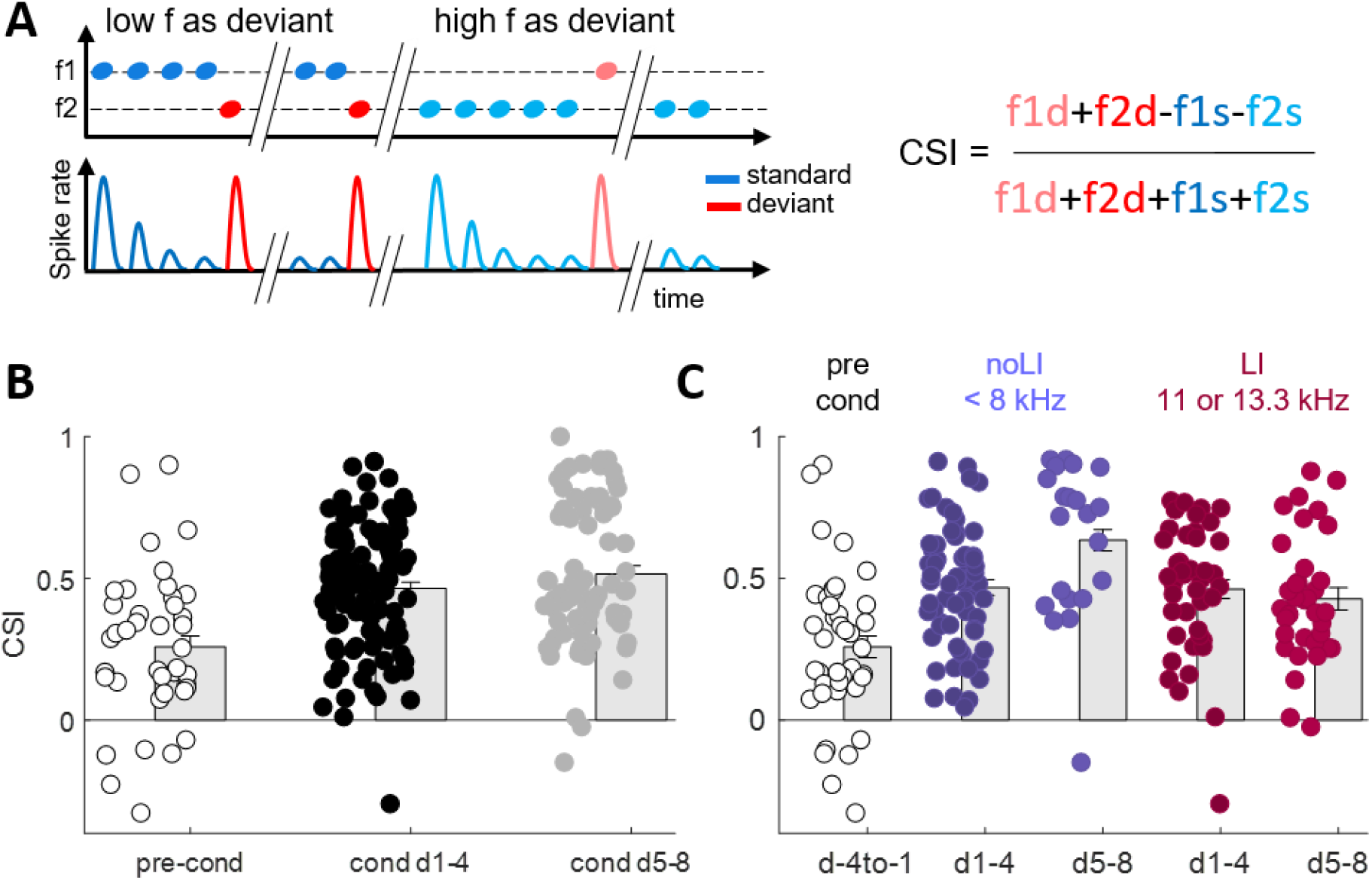
stimulus-specific adaptation was increased in conditioned mice. **A. left:** Schematic of odd ball paradigm. Low frequency (f2) was first played as deviant (red) against f1 (blue), then as standard (light blue) against f1 as deviant (light red); **right:** the combined stimulus-specific adaptation index (CSI) was calculated as the normalized mean of the difference between all deviant responses and all standard responses. **B.** CSI values across recording sites during preconditioning (white; n=7 mice and 41 recordings), first 4 conditioning days (black; n=18 mice and 106 recordings), and days 5 to 8 of conditioning (gray; n=17 mice and 76 recordings). **C.** Mean CSI according to exposure frequency (color coded) and time-block of conditioning (left: days 1 to 4, or right: days 5 to 8). Preconditioning (white; n=7 mice and 41 recordings), exposed to tones < 8 kHz (purple; conditioning d1-4 n=9 mice and 63 recordings; d5-8 n=9 mice and 41 recordings), exposed to 11.2 or 13.3 kHz (red; conditioning d1-4 n=9 mice and 43 recordings; d5-8 n=8 mice and 35 recordings). f: frequency.

### Recording selection and statistics

In each mouse, recordings were taken from 1 to 3 recordings sites. In each recording site, the electrode was lowered perpendicular to dura over the auditory cortex to a depth of 350 to 450 μm (layer 4) and 1 or 2 pairs of frequencies were played in oddball paradigms. Responses to each frequency were selected and included based on a significant response during the deviant-alone protocol. Response rate was calculated based on the number of spikes in an interval of 100 ms starting at stimulus onset. Common SSA indices were used to quantify the level of SSA (See Figure 4A; CSI=(deviant-standard)/(deviant+standard), where deviant and standard represent the average number of spikes in deviant and in standard trials, respectively, summed over the two frequencies used in the test).

In order to take the potential similarity of responses within mouse into account, we used extensively linear mixed effects models with mouse-dependent random factors. The models were fitted using the matlab function fitlme (Matlab 2020a, Mathworks). In a linear mixed effects model, all effects are linear (as in ANOVA or in general linear models). The effects are either ‘fixed’ or ‘random’. Fixed effects are estimated as free coefficients (again, similar to standard ANOVA or general linear models), while random effects are assumed to be Gaussian with mean 0 and a covariance matrix that is estimated from the data by maximum likelihood. In the models used here, the neural responses of each unit to the two frequencies tested in the oddball sequences could enter into the analysis. These responses are however highly correlated. To account for this correlation, a unit-specific random effect was used for the two frequencies. This random effect is different for each unit at that experimental time point and is estimated under the assumption that it has a Gaussian distribution with mean zero. The similarity between the responses of the same unit to the two frequencies is therefore captured by the variance of this random effect and is used as an additional source of variation when estimating significance.

## Results

To better understand how previous experience influences learning-driven plasticity in sensory gating in the primary auditory cortex (A1), we sampled the activity of neurons in mice that had been exposed to different frequencies in an automated behavioural apparatus, the Audiobox (Figure 1A), before being conditioned to a 13.3 kHz tone.

During a pre-conditioning phase, mice living in groups of 8-10 individuals in the Audiobox heard either of 2 pure tones in each visit to the water corner. In 83% of the visits, mice heard a 6.7 kHz tone and in 17% of the visits, a mouse-specific frequency (pre-conditioning exposure frequency of 3.3, 5.6, 7.9, 11.2 or 13.3 kHz; Figure 1C). All visits in this phase were ‘safe’ and water was available irrespective of the tone frequency (Figure 1A-B, methods). Mice nose-poked in most visits, avoiding a nose-poke in about 30% of visits (Figure 1D, days −4 to −1).

Following pre-exposure, during the conditioning phase, mice in all groups heard the same tones. The safe 6.7 kHz tone was still presented in 83% of the visits as before. In the remaining 17% of the visits, now ‘conditioned’ visits, a 13.3 kHz tone was presented to all animals in all groups (Figure 1B, bottom). Importantly, the 13.3 kHz tone was novel to all the mice except those that had heard it during pre-exposure. A nose-poke in the conditioned visits was followed by an air puff and access to water was denied.

As already reported (de Hoz and Nelken 2014), the mice learned to avoid nose-poking in conditioned visits at a rate that depended on the pre-conditioning frequency (Figure 1D, circled red). An ANOVA on percentage nosepoke avoidance per day with visit sound type (6.7 kHz −83% of visits-vs rare sound −17% of visits-), phase (pre-exposure vs conditioning) and pre-exposure frequency as factors, revealed a strong effect of sound type (F(1,678)=242.08; p<0.0000), and phase (F(1,678)=834.16; p<0.0000), and an almost significant effect of frequency (F(4,678)=2.36; p=0.0519), as well as interactions between sound type and phase (F(1,678)=427.05; p<0.0000), and phase and frequency (F(1,678)=4.46; p=0.0014). Focusing on the conditioning phase, an ANOVA with sound type, day of conditioning, and pre-exposure frequency revealed an effect of sound type (F(1,274)=543.48; p<0.0000), and frequency (F(4,274)=4.86; p=0.0008), a strong sound type by day interaction (F(5,274)=16.21; p<0.0000) and a sound type by frequency interaction (F(4,274)=3.49; p=0.0084). Mice that had been exposed to frequencies below 8 kHz (3.3, 5.6 and 7.9 kHz) and, therefore, sufficiently different from the conditioned frequency, learned to avoid nose-poking in conditioned visits very fast. This is evident in the rapid increase in avoidance within the first 12 hours of conditioning (Figure 1E, purple, blue and gold). The learning rate was comparable to that of mice that have not been pre-exposed, who often learned to avoid noise-poking to the conditioned frequency within a single trial (de Hoz and Nelken 2014; Chen, Krueger-Burg, and de Hoz 2019).

Mice exposed to 11.2 and 13.3 kHz, on the other hand, presumably surprised by the sudden change of valence of the pre-exposure tone, nose-poked more persistently during the initial conditioned visits (Figure 1D, circled red; and Figure E, orange and red), an effect described as latent inhibition and thoroughly characterized in de Hoz & Nelken (2014; cf. Figure 5). Indeed, an ANOVA on day 1 of conditioning, with sound type and pre-exposure frequency as factors, revealed a strong effect of sound type (F(1,72)=36.99; p<0.0000), no effect of frequency (F(4,72)=1.87; p=0.1244), and a strong interaction (F(4,72)=5.47; p=0.0007), such that when focusing on the conditioning sound a strong effect of pre-exposure frequency was observed (F(4,36)=3.7; p=0.0126). This effect was short lasting and, after 18 hours, all mice had reached asymptotic levels of avoidance when presented with the conditioned tone (Figure 1E, orange and red). Thus, exposure to a given frequency generated an expectation about the valence of this frequency that extended to nearby frequencies and was unveiled through latent inhibition (LI).

In addition to the avoidance of the conditioned tone, all groups showed an increase in the avoidance during ‘safe’ visits (Figure 1D, green, note increase in avoidance with respect to pre-conditioning levels in all groups). This generalization of avoidance to the safe sound depended on the pre-exposure frequency on day 1, much like the avoidance of the conditioned tone (F(4,36)=3.66,p=0.0134), was more pronounced in mice exposed to 11.2 and 13.3 kHz and lasted longer than the latent inhibition.

Indeed while in mice exposed to 11.2 and 13.3 kHz, the avoidance in safe visits decreased across days of conditioning (F(7,90)=5.64,p<0.0000), it started lower and did not change in mice exposed to frequencies below 8 kHz (F(5.68)=1.16,p=0.3363). Thus, by the time the mice consistently avoided nose-poking in conditioned visits (Figure 1D, red circles) they still showed a higher rate of avoidance of the safe tones. Because of the differential effect of pre-exposure to frequencies near and far from the conditioned tone, mice pre-exposed to frequencies below 8 kHz were analyzed together (these were animals that didn’t show much latent inhibition in the behavioral tests, denoted below as noLI), as were animals pre-exposed to frequencies above 8 kHz (11.2 and 13.3 kHz; these animals showed strong latent inhibition, denoted as LI).

The aim of this study was to understand the effect that sound experience had on auditory responses in auditory cortex (AC), and in particular on the processes tagged by SSA – the adaptation of the responses to the standard tone and the release from adaptation during presentations of the deviant tones (Figure 2A, bottom half; blue represent standard tones and red represents deviant tones). Single mice were removed from the Audiobox at different time points before and after conditioning (Figure 2A) for acute electrophysiology. We presented different tone pairs (ΔF of 20-40%) in an oddball sequence, where a sequence of tones with a common frequency (standard, 95% of trials, 475 repetitions) was randomly interrupted by tones of a different frequency (deviant, 5% of trials, 25 repetitions; Figure 2A, middle).

We first measured adaptation to the standard tones at three time points along the time course of training. To capture the slow time course of avoidance of the safe tone (that disappeared earlier in the noLI group than in the LI group), we binned the experimental time to pre-conditioning (days −4 to −1, before conditioning began), early conditioning (days 1-4), during which both LI and noLI animals showed avoidance of the safe tone, and late conditioning (days 5-8), when the LI animals still showed an elevated level of avoidance of the safe tone while noLI animals already returned to largely normal level of avoidance of the safe tone. Figure 2B shows the average responses (over all tested frequencies and all pre-exposure groups) of the standards as a function of their position in the sequence of 475 repetitions, separately for each of the three experimental time points. The plot suggests that the initial, unadapted, responses (at the beginning of the sequence) tended to be stronger after conditioning (darker circles above the open circle). In contrast, the adaptation that ensued upon repetition of the standard tone was relatively mild before conditioning, but was deeper following conditioning. In consequence, the responses recorded in conditioned animals towards the end of the sequence (Figure 2B, filled circles) were smaller than the responses in animals before conditioning (open circles).

We fitted a linear mixed-effects (LME) model to the data (209475 individual responses from 223 units recorded in 44 mice). The experimental time point (before conditioning, days 1-4 and day 5-8) and the inverse of the trial number within the block (Antunes et al., 2010 PMID: 21124913) were used as fixed factors. Unit-dependent random intercepts, separately for each experimental time point, were used for test frequency (high/low).

There was a large effect of trial number (F(1,209469)=83.5,p=6.4e-20), in consequence of the large reduction in responses along the sequence of trials. While the main effect of experimental time point was only weakly significant (F(1,209469)=3.40,p=0.033), there was a significant interaction between trial number and experimental time point (F(2,209469)=34.5, p=1.0e-15), reflecting the inversion in the response size between early and late responses before and after conditioning. Because of the significant interaction, we verified the effect of experimental time point separately for the first standard trial in each block and for the average of the last 75 standards in each block. Both were significant (first trial, effect of time point: f(2,438)=4.03, p=0.018; last trials, effect of time point: F(2,44097)=4.96, p=0.0070). However, the direction of the effects was reversed: the first standard responses were larger on average after conditioning than before conditioning, while the last standard responses were smaller on average after conditioning than before conditioning.

To capture the time course of adaptation within blocks, we used the responses to the 1^st^, 5^th^ and 20^th^ standard trial in each block, as well as the average responses of the last 75 trials (28158 individual responses in 182 units recorded in 36 mice). These trials represented well the trajectory of the adaptation. The data are separated in a 2×2 design by experimental time point and by pre-exposure frequency binned to frequencies below 8 kHz (noLI group) and frequencies above 8 kHz (LI group).

Figure 2C shows the main effects of interest in this study. It suggests that the adaptation time course (as captured by the responses at the corresponding trial numbers) differed between the LI and noLI animals in ways that depended on the time after conditioning. To analyze the data, we used a linear mixed effects model. The three fixed effects were as described above the trial number within block (4 levels), the pre-exposure frequency (2 levels, noLI and LI) and the experimental time point (2 levels, 1-4 and 5-8 days after conditioning). All pairwise interactions were included in the model as well. As before, unit-dependent random intercepts were used for test frequency (low or high), separately for each experimental time point.

The strong adaptation within sequences was reflected, as expected, by a highly significant main effect of trial number (F(3,28140)=125, p=2.1e-80). There was also a strong main effect of pre-exposure frequency (F(1,28145)=32.0,p=1.5e-8), with the LI animals (pre-exposed to 11 or 13 kHz, blue hues in Fig. 2C) showing on average smaller responses than the noLI animals (pre-exposed to frequencies <8 kHz, red hues in Fig. 2C). Both of these main effects are clearly apparent in Fig. 2C.

The effect of exposure frequency (noLI vs. LI animals) was modulated by a significant interaction between exposure frequency and trial number (F(3,28145)=12.9, p=2.1e-8), due to the different effects of pre-exposure frequency on the time course of adaptation to standards. Indeed, while the responses at the first standard trial, 5^th^ trial (marked with larger dots for visualization in Figure 2C), and 20^th^ trial were on average significantly smaller in LI animals than in noLI animals (F(1,359)=4.33, p=0.038; F(1,359)=5.18, p=0.023; F(1,359)=4.18, p=0.042 respectively), there was no significant difference between the average responses of LI and noLI animals for the last standard trials in a block (F(1,359)=0.539, p=0.46). Thus, on average in a given trial sequence, the noLI animals had larger initial responses than LI animals, but eventually both groups showed adaptation to similar activity levels (lower after conditioning than before, as shown above).

There was no main effect of the experimental time point (days 1-4 vs. days 5-8 after conditioning, F(1,28140)=0.101, p=0.75). However, the experimental time point did affect the responses through two interactions. First, the adaptation patterns differed significantly between animals tested in days 1-4 vs. days 5-8 after conditioning. Indeed, there was a significant interaction between time point and trial number (F(3,28145)=4.18, p=0.0057), driven by a slower adaptation on days 5-8 after conditioning than in days 1-4 after conditioning, and effect reflected mainly in the larger responses to trial 5 in the data from animals tested 5-8 days after conditioning.

Most importantly, there was a significant effect of experimental time point for mice showing latent inhibition). This interaction was due to the increase of all the responses of LI animals (but not of noLI animals) tested on days 5-8 after conditioning relative to animals tested on days 1-4 after conditioning.

In summary, in both LI and noLI animals, conditioning increased the unadapted responses to sound (first standards) but also increased the depth of the adaptation, resulting in smaller response levels for standards at the end of the sequence. Neurons in noLI animals showed the same adaptation pattern at the two experimental time points, while neurons in LI animals showed additional changes in the responses between the two experimental time points: responses to standards in the first experimental time point were smaller than in noLI animals and adaptation was faster, while in the second experimental time point responses increased and the adaptation slowed down so that the responses matched more closely those of neurons recorded in noLI animals.

The deviant responses were also sensitive to conditioning. Figure 3A shows the average responses to deviant tones as a function of their sequential number along the sequence, for the three time points (before conditioning, days 1-4 after conditioning, and days 5-8 after conditioning). For the statistical analysis, we used the same model as for the standards. The main effect of trial number was not significant (F(1,11019)=3.61, p=0.058), reflecting the almost constant responses to deviants along the oddball sequence. There were however a significant main effect of experimental time point (F(2,11019)=5.71, p=0.0033) reflecting the larger responses to deviants after conditioning than before conditioning. There was also an interaction between experimental time point and trial number (F(2,11019)=3.25, p=0.039), reflecting the small increase in the first few deviant responses in animals before conditioning, which was absent from the average response after conditioning.

We then averaged the responses to the 25 deviant presentation within each block and analyzed them as a function of experimental time point (d1-4 or d5-8) and pre-exposure frequency (noLI vs. LI). Since the interaction was not significant, we fitted a main effect model to the data. As expected, there was a significant effect of experimental time point (days 1-4 after conditioning vs. days 5-8 after conditioning; F(1,9022)=5.83, p=0.016), with deviant responses recorded on days 5-8 being on average larger than on days 1-4 after conditioning.

The main effect of pre-exposure frequency on deviant responses was not significant (F(1,9022)=1.66, p=0.20).

The interaction between adaptation to the standard sound and maintained sensitivity to the deviant is typically measured using the contrast between the deviant and the standard responses to the same tone (f1 or f2, Figure 4A). The combined stimulus-specific adaptation index (CSI) for both f1 and f2 is calculated as described in Figure 4B. Since conditioning resulted in a deeper adaptation with a small increase of the deviant responses, the contrast between deviant and standard responses increased after conditioning (Figure 4B; main effect of experimental time point, F(2,215)=18.6, p=3.4e-8; 218 individual CSI values recorded in 44 mice; linear mixed effects model with random intercepts for mice nested within experimental time points).

We tested the effect of pre-exposure frequency on the responses after conditioning by testing the dependence of CSI on the experimental time point and the pre-exposure frequency. The main effect of pre-exposure frequency was not significant (F(1,175)=0.0154, p=0.90), but there was a significant main effect of experimental time point (F(1,175)=7.42, p=0.007) as well as a significant interaction between pre-exposure frequency and experimental time point (F(1,175)=5.41, p=0.021).

To find the origin of the interaction, we tested separately the effect of experimental time point in the LI and the noLI animals. The effect of experimental time point was significant in the noLI animals (F(1,99)=7.03, p=0.0093), with the CSI being on average larger in units recorded 5-8 days after conditioning, but was not significant in the LI animals (F(1,76)=0.467, p=0.50), with the CSI being smaller (although not significantly so) in units recorded 5-8 days after conditioning.

To conclude, paralleling the behavioral pattern, changes in sound evoked responses in A1 also combined effects at different time scales. Overall, unadapted evoked activity (both to first standards and to deviant tones) increased following conditioning, but this increase developed more slowly in the LI group. Adapted responses, on the other hand, were weaker in conditioned mice. This effect was particularly large at the LI group in the first experimental time point (1-4 days after conditioning). The effects were independent of the frequency tested (data not shown).

## Discussion

In this study, we set out to quantify the effect of prior sound experience on conditioning-triggered changes in the dynamics of neuronal responses. We have shown before (de Hoz and Nelken 2014) that prior exposure to frequencies near the conditioned tone led to behavioral latent inhibition – a slowing down of the acquisition of the conditioned response (LI-groups). The delay in conditioning lasted less than 24 hours and all groups showed comparable levels of avoidance of the warning sound by day 2 of conditioning. In contrast, the generalization of avoidance to the familiar safe tone was more persistent and lasted over 4 days in the LI groups. Groups exposed to tones away from the conditioned frequency showed neither latent inhibition (noLI groups) nor a long lasting (>4 days) generalization to the safe sound.

Here we report that in parallel with the behavioral effects, electrophysiological responses were influenced by both prior exposure and conditioning. In particular, prior exposure led to remarkably long-lasting electrophysiological consequences that depended on the pre-exposure frequency. Behaviorally, latent inhibition was relatively short lasting, observed mainly in the first 6 hours of conditioning to the 13.3 kHz tone, when the mean performance of mice exposed to frequencies below 8 kHz was higher than that of mice exposed to 11.2 or 13.3 kHz. Since we began collecting electrophysiological data only 24 hours after the beginning of conditioning, this study lacks the resolution to discern plastic changes within the first few hours following conditioning. On the other hand, the time scale of the electrophysiological data corresponds to the changes we observed in the excess avoidance of the safe tone.

Conditioning led to a general increase in unadapted sound evoked responses, reflected in the strong responses to the first standard tone in all conditioned groups and at all time points, as well as in the increase in deviant responses. Such changes are expected (Engineer et al. 2004).

Conditioning also led to an increase in the depth of adaptation to repeated standards. In consequence of these changes, the contrast between standards and deviants (quantified as CSI) was larger after conditioning than before. This result is somewhat different than that in a previous report of plasticity of SSA (Yaron et al. 2020), where the responses to pure tones used in conditioning and then tested as standards increased overall (rather than decreasing, as here). Potential reasons for this difference may be species differences (rats in Yaron et al. 2020, mice in the current report), and the use of anesthesia (Yaron et al. recorded in awake animals while the test recording in the current report were performed in anesthetized animals), but potentially the most important difference is the learning paradigm – Yaron et al. used classical fear conditioning, while the current report used operant conditioning.

Similar increases in both sensory responses and depth of adaptation have been reported earlier in animals exposed to enriched environments with predictable sounds (Engineer et al. 2004). Thus, the larger initial responses and the deeper adaptation may be a consequence of sound processing in environments that contain predictable sounds (Cruces-Solís et al. 2018).

The most important finding of this paper is the long-term dynamics of the plasticity in the LI animals. These slow plastic changes paralleled the slow changes in behavior that we have documented previously (de Hoz and Nelken 2014) and Fig. 1). Thus, exposure in an enriched environment, when followed by an incongruous event (the conditioning of a previously safe tone, LI mice) leads to long lasting changes in behavior, reflected in the very slow relearning that 6 kHz is safe, as well as long lasting changes in the adaptation profile. Both are different to those of animals exposed and conditioned to different frequencies (noLI mice).

Indeed, following conditioning, the responses evolved differentially in noLI and LI animals over the experimental time scale. While in noLI animals the adaptation pattern was roughly stable over time, in LI animals the responses to standards on days 1-4 were weaker than in the noLI animals (Fig. 2C), but increased on days 5-8, approaching the pattern of the electrophysiological responses in the noLI animals. Since the behavior of the LI animals also approached that of the noLI animals on days 5-8, the convergence of the electrophysiological responses tracked the convergence in behavior.

If so, it may be that the difference in electrophysiological responses on days 1-4 between the noLI and the LI animals is the factor that drives the higher level of avoidance of the safe tone in the LI vs. noLI animals. We recognize that the avoidance of the safe tone may be due to multiple mechanisms, only part of them directly related to the auditory processing of the incoming stimuli. For example, In the following, we discuss potential auditory processing mechanisms based on the responses that we measured that may link the changes in neural responses reported here with the changes in behavior.

The auditory information that feeds into the behavioral choices is most likely determined by the responses to the first few sounds in the Audiobox. Indeed, the decision to nose-poke or not is typically made within the first 5 seconds, with only a few sound presentations before decision (de Hoz and Nelken 2014). The responses evoked by these first few stimuli correspond best with the responses to the first few sound presentations of the oddball sequence. Under these conditions, more robust responses would make the neural discrimination task easier, because the difference between the responding and non-responding populations would be larger and easier to detect.

In order to perform the decision, it is necessary to compare the responses of the neuronal populations responding to the two tones (for example by subtracting or dividing the total spike counts evoked by the two tones – but the exact operation is not important here), and to select a threshold on the result. Our behavioral observations (Fig. 1) show that animals select this discrimination threshold such that avoidance of the conditioned tone is very high. However, because of neural noise, any threshold would result in a certain level of misdetection of the safe tone (and therefore avoidance of the safe tone). For the same level of avoidance of the conditioned tone, the level of avoidance of the safe tone would depend on how discriminable the responses are. More robust neural responses (to either tone) would result in larger differences between the responses of the populations responding to the two tones and therefore to lower level of avoidance of the safe tone.

The crux of the argument is the observation that on days 1-4, neurons in noLI animals showed larger responses to the initial tones in the oddball sequences than neurons in LI animals. On days 5-8, the large responses in the noLI animals remained and while the responses in the LI animals increased, tending towards those in noLI animals. Assuming the same discrimination mechanism in the two groups, including the setting of the threshold, the argument above suggests that on days 1-4 the LI animals would show larger avoidance of the safe tone than the noLI animals, and that this difference would decrease in days 5-8. This is indeed the behavior we observed.

While this argument accounts for the difference between the noLI and the LI animals, it doesn’t account for the decline of the avoidance of the safe tone in the noLI animals. Using similar arguments, this decline may be related to a slow shift of the threshold of discrimination. The full explanation has therefore two components: slow shift of the decision threshold which is common to the two groups in parallel with the improvement in signal-to-noise ratio of the neuronal responses in the LI animals over the same period of time.

This interpretation has an obvious correlate – trials in which mice avoid the safe tone are also trials in which the resulting responses in auditory cortex are weaker. Experiments in which behavior and neural responses are measured simultaneously will be necessary in order to check this hypothesis.

Numerous studies have contributed to the characterization of SSA (Harpaz et al. 2021; Carbajal and Malmierca 2018) but very little is known about how it is affected by experience. In a previous study (Yaron et al. 2020), we found that rats trained to discriminate between two words, showed increased SSA to words and tones, consistent with our data. In the current study, unlike in Yaron et al. (2020), electrophysiology was performed under anaesthesia. While SSA is largely the same with and without anesthesia (Polterovich, Jankowski, and Nelken 2018), anesthesia dampens inhibition (Guo et al. 2021)and might have diminished frequency-specific effects on response sensitivity.

Overall our data reveal that slow changes in behaviour that accompanied conditioning, such as the generalization of conditioning to a safe tone, paralleled the slow dynamics of experience-dependent plasticity in auditory cortex. Prior experience had both fast and slow effects on behavioural responses that were paralleled by differential dynamics of stimulus sensitivity and adaptation.

